# Improving the reliability of T_2_ measurement in magnetic resonance imaging

**DOI:** 10.1101/2024.06.09.598128

**Authors:** Xiuli Yang

## Abstract

Magnetic resonance imaging (MRI) is a versatile technique broadly utilized in research and clinic. Among the information available from MRI measurements, transversal relaxation time (T_2_) is a basic but critical one to reflect the microstructure and microenvironment at the molecular level. A popular method to estimate T_2_ is employing a series of evolution time (TE) values and, thereafter, fitting with the exponential model (termed as T_2_ fitting). Despite of the popularity in using T_2_ fitting, there is a lack of knowledge how related experimental parameters, such as signal-to-noise (SNR), number of TE utilized, dynamic range of TE, and repetition number of each TE, affect the measurement reproducibility. In this study, we performed Monte Carlo simulations to investigate how SNR, TE number, TE range, and repetition number influence the reliability of T_2_ measurement, which was indexed by coefficient of variations. Good reliability with ≤5.0% variation can be achieved when the normalized noise level is below 2.2%. Optimal TE range for measuring T_2_ is related to the T_2_ under evaluation. TE number and repetition number can be increased to reduce measurement variations.

## Introduction

Since its advent, magnetic resonance (MR) technique has been extensively developed and broadly utilized in different research and industrial fields (*1-5*). The versatility in adapting pulse sequences for different spatial (*6, 7*) or spectral information (*8, 9*) is an outstanding feature adding the popularity and irreplaceability of MR technique. Among the abundant information from MRI, transverse relaxation time (denoted as T_2_) is a most basic but critical one because T_2_ is sensitive to the microstructural and microenvironmental changes (*10-13*). For example, in the hemorrhagic stroke, with the development of edema, there will be distinct increase in T_2_ due to the accumulation of mass free water around the vascular rupture (*14*). T_2_ is not only sensitive to acute pathogenesis as in the strokes, but also indicative for chronic pathological development, e.g., in the Alzheimer’s disease (*15*). Systematic development of amyloid plaque and neurofibrillary tangles can change the microenvironment around neuronal cells and become observable via the T_2_ monitoring (*16, 17*).

One most inspirational application of T_2_ measurement lies within the functional MRI (fMRI) study. There was a well-known relationship between blood oxygenation and blood T_2_ (*18, 19*). Based on this calibratable relationship, novel MRI techniques have been developed to unravel an important parameter related to brain metabolism, namely, oxygen extraction fraction (OEF) (*20-23*). Along this direction, cerebral blood flow (CBF) (*24, 25*) can be measured to estimate the cerebral metabolic rate of oxygen (CMRO_2_) (*26, 27*), which is direct measurement on the level of aerobic metabolism in brain, using the Fick’s principle. Therefore, T_2_ is associated with rich physiological information and provides valuable opportunities to unravel the brain functions.

The transversal relaxation follows exponential decay and quantitative T_2_analyses can be achieved by fitting data with the exponential model. To introduce different weighting, a series of evolution time (denoted as TE) is often employed. Critical parameters relevant to the T_2_ fitting include the noise level, number of TE, range of TE, and repetition number of each TE. The effects of these experimental parameters on the reliability of T_2_ measurement have not been systematically examined in the literature. Therefore, we aim at filling this knowledge gap with the present study. Note that the time cost associated with an experiment study exploring the influences of experimental parameters will be prohibitively high. As an alternative, we employ the Monte Carlo simulation method to explore the dependence of measurement reliability on the basic experimental parameters with dense acquisitions.

## Methods

### Monte Carlo Simulation

Monte Carlo simulation was performed to investigate the relationship between measurement reproducibility of T_2_ and experimental parameters. Critical parameters in T_2_ mapping consisted of the noise level, dynamic range of TE, number of TE, and number of repetitions. Four sub-studies were performed to determine the influences of these experimental parameters on the reproducibility of T_2_ measurements. Simulations were performed with 2k repetitions to evaluate the coefficient-of-variation (CoV). All scripts were written based on the MATLAB platform.

#### Study 1: Dependence of T_2_ measurement on noise levels

Five different T_2_ values were simulated, including 15, 20, 25, 30, and 35 ms. Normalized noise levels by reference to the signal intensity at TE=0 ms were used in this study. Fifty noise levels ranging from 0.1% to 5.0% at the interval of 0.1% were utilized. TE range was twice the ground-truth T_2_ value, e.g., TE range of 30 ms was used when measuring the ground-truth T_2_ of 15 ms. A total number of 10 TE values were utilized.

#### Study 2: Dependence of T_2_ measurement on the dynamic range of TE

Five different T_2_ values were simulated, including 15, 20, 25, 30, and 35 ms. The noise level of 2.2% was considered. Fifty TE ranges from half to 10-fold of the ground-truth T_2_ value were investigated. A total number of 50 TE values were utilized.

#### Study 3: Dependence of T_2_ measurement on the total number of TE

Five different T_2_ values were simulated, including 15, 20, 25, 30, and 35 ms. The noise level of 2.2% was considered. The TE range of 1.857 ground-truth T_2_ value was employed. The TE number from 3 to 30 was examined for comparisons. Exact TE values were equidistant and lied within 0 and TE range.

#### Study 4: Dependence of T_2_ measurement on the repetition number

Five different T_2_ values were simulated, including 15, 20, 25, 30, and 35 ms. The noise level of 2.2% was considered. The TE range of 1.857 ground-truth T_2_ value was employed with 5 equidistant TE. Repetition numbers from 1 to 20 were compared.

### Statistical analyses

Linear regression model was utilized to explore the influence of different experimental parameters on the reliability of T_2_ measurements. A P<0.05 was considered significant.

## Results

### Study 1: Dependence of T_2_ measurement on noise levels

Based on the linear regression analyses, the standard deviation of fitted T_2_ exhibited a significant positive dependence on the noise level (P<0.0001), but the averaged fitted T_2_ did not depend on the noise level (P=0.20), suggesting that the fitted T_2_ did not suffer from dramatic inaccuracy when sufficient averages were available (Figure 1A). Interestingly, the standard deviation of fitted T_2_ was associated with a significant positive interaction effect, i.e., T_2_-by-noise-level, effect (P<0.0001), suggesting that standard deviation was larger when measuring higher T_2_. Coefficient of variation (CoV), which is an index used to assess the reproducibility, exhibited significant dependences on noise level (coefficient = 2.127; P<0.0001) and T_2_ values (coefficient = -0.010; P<0.0001) (Figure 1B). Over the tested range (i.e., 15-35 ms), the T_2_ effect on CoV will lead to a CoV difference of 0.2%, which is a negligible effect in comparison with the noise effect on CoV. If a CoV of 5.0% is allowed, the cut-off normalized noise levels were 2.2%, 2.5%, 2.4%, 2.3%, and 2.4% for measuring T_2_ of 15 ms, 20 ms, 25 ms, 30 ms, and 35 ms, respectively. Thereafter, the normalized noise level at 2.2% is used to ensure that variations are within 5.0% across different ground-truth T_2_s.

**Figure 1.**
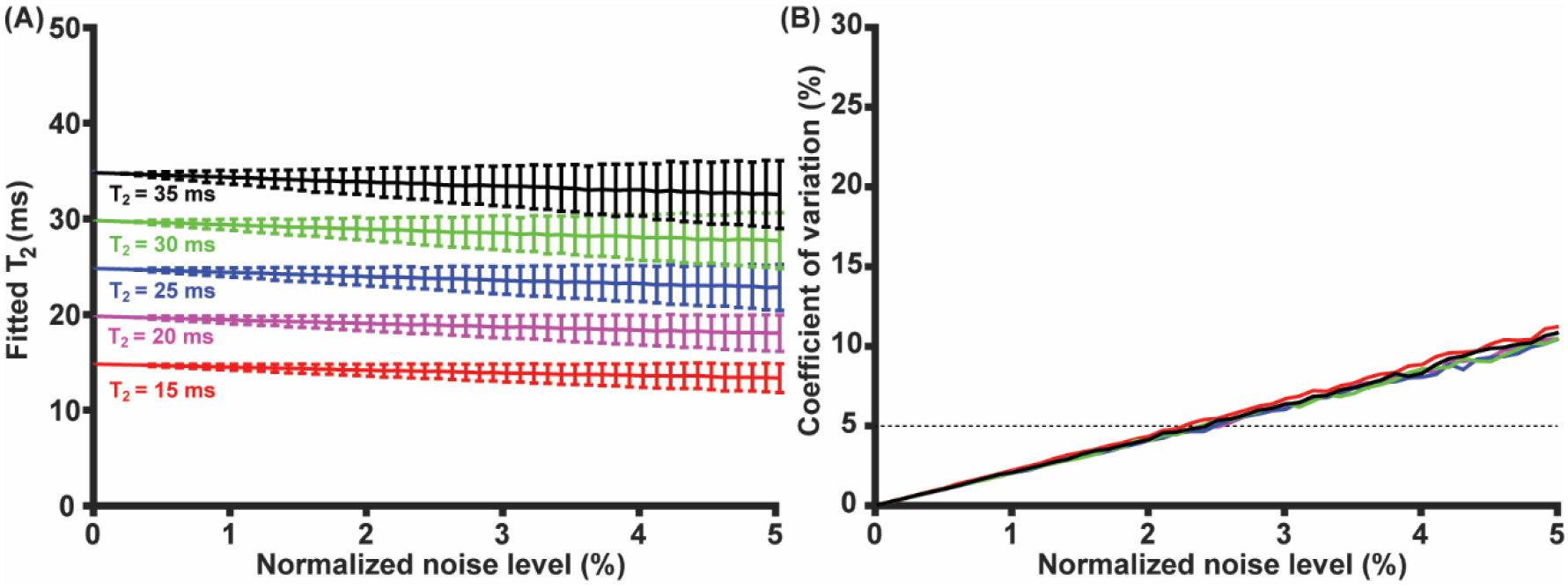
Dependence of measured T_2_ (A) and CoV (B) on the normalized noise level. Error bar stands for the standard deviation.

### Study 2: Dependence of T_2_ measurement on the dynamic range of TE

The CoV of T_2_ measurement depended on the normalized TE range (coefficient >0, P<0.0001) but not the ground-truth T_2_ values (P=0.99) (Figure 2A). Note that the dependence of CoV on normalized TE range was nonlinear (Normalized-TE-Range^2^: P<0.0001). We further fitted the CoV curves according to polynomial functions with different orders. It can be observed that R^2^ was approaching 1.0 with increased orders. Using 0.99 as the threshold value for R^2^, the polynomial fitting order was ≥7 (Figure 2B). At the 7-order fitting, we estimated that the minimal CoV arose at the normalized TE range of 1.857. These results indicate that the optimal TE range for T_2_ measurement is 1.857T_2_, where T_2_ is the transversal relaxation time to be measured.

**Figure 2.**
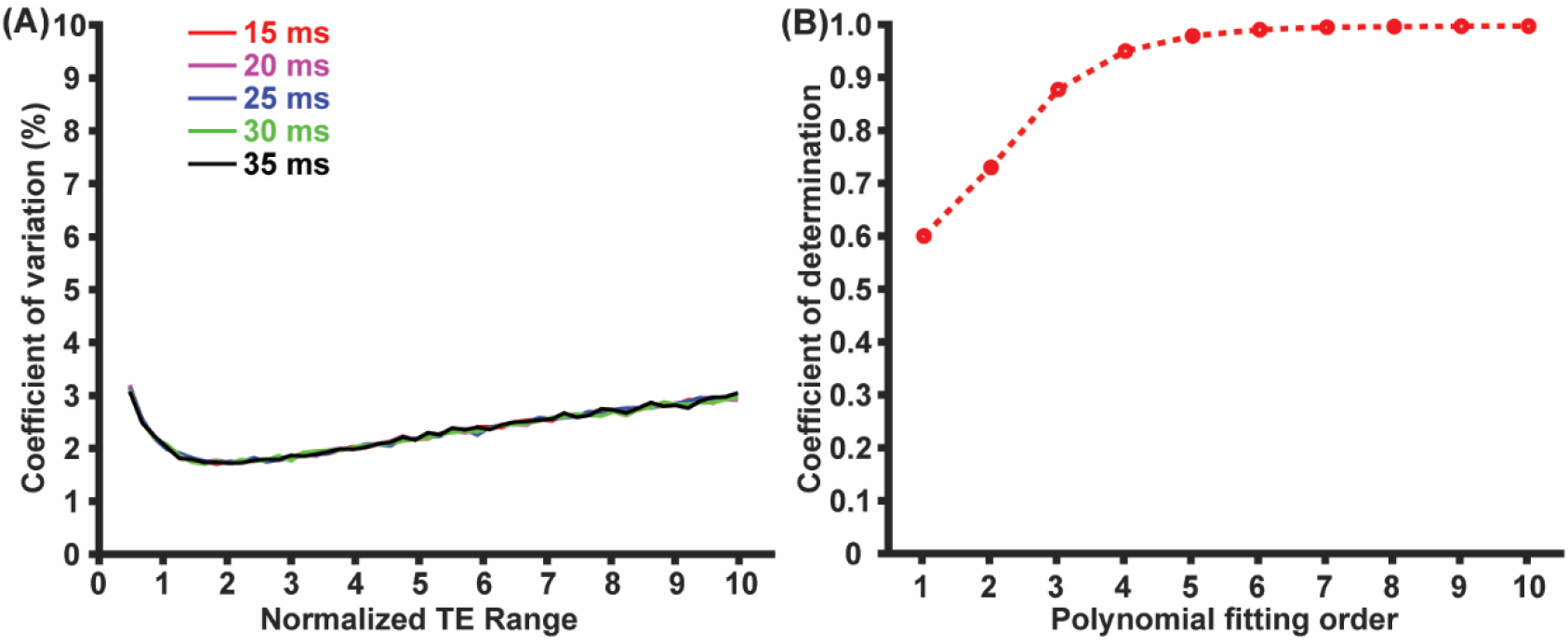
CoV as functions of normalized TE range (A) and coefficient of determination (i.e., R^2^) as a function of the order of polynomial fitting order.

### Study 3: Dependence of T_2_ measurement on the total number of TE

The CoV of T_2_ measurement exhibited a significant dependence on the TE number (coefficient < 0, P<0.0001) but not on the ground-truth T2 values (P=0.99) (Figure 3A), suggesting that variations of T_2_ measurement can be alleviated by adding the number of TE values. The TE-number effect was nonlinear (TE-number^2^ term, P<0.0001). We further fitted the CoV curves with polynomial fitting and found that R^2^≥0.99 can be obtained with polynomial fitting ≥4 orders (Figure 3B). With the 4-order fitting, we estimated that CoV<5% could be obtained with TE number ≥5.

**Figure 3.**
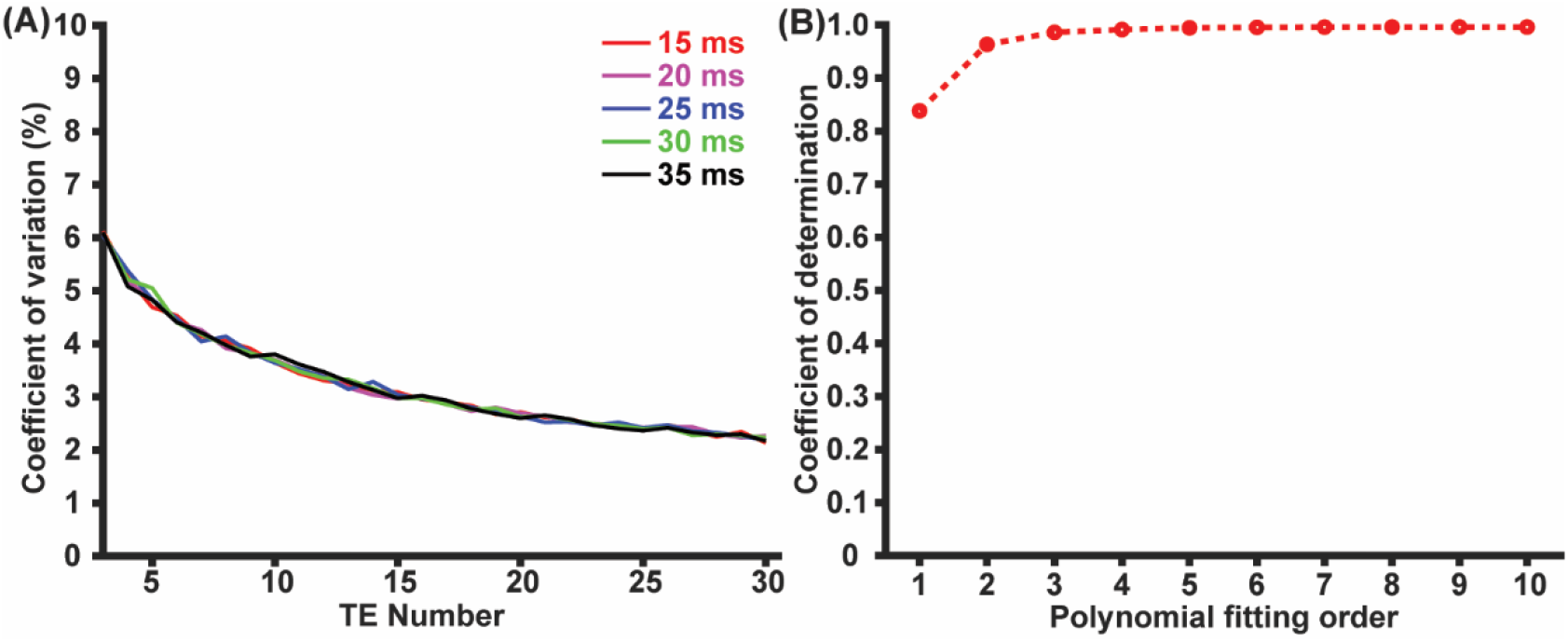
CoV as functions of TE number (A) and coefficient of determination as a function of polynomial fitting order.

### Study 4: Dependence of T_2_ measurement on the repetition number

The CoV of T_2_ measurement depended on the repetition number (coefficient < 0, P<0.0001) but not on the ground-truth T_2_ values (P=0.79) (Figure 4A), suggesting that adding the repetition number can help alleviate the variation and thereafter improve reproducibility. The dependence of CoV on repetition number was nonlinear (Repetition-number^2^ effect, P<0.0001). Polynomial fittings were performed to the CoV data and R^2^≥0.99 was obtained with fitting ≥5 orders (Figure 4B).

**Figure 4.**
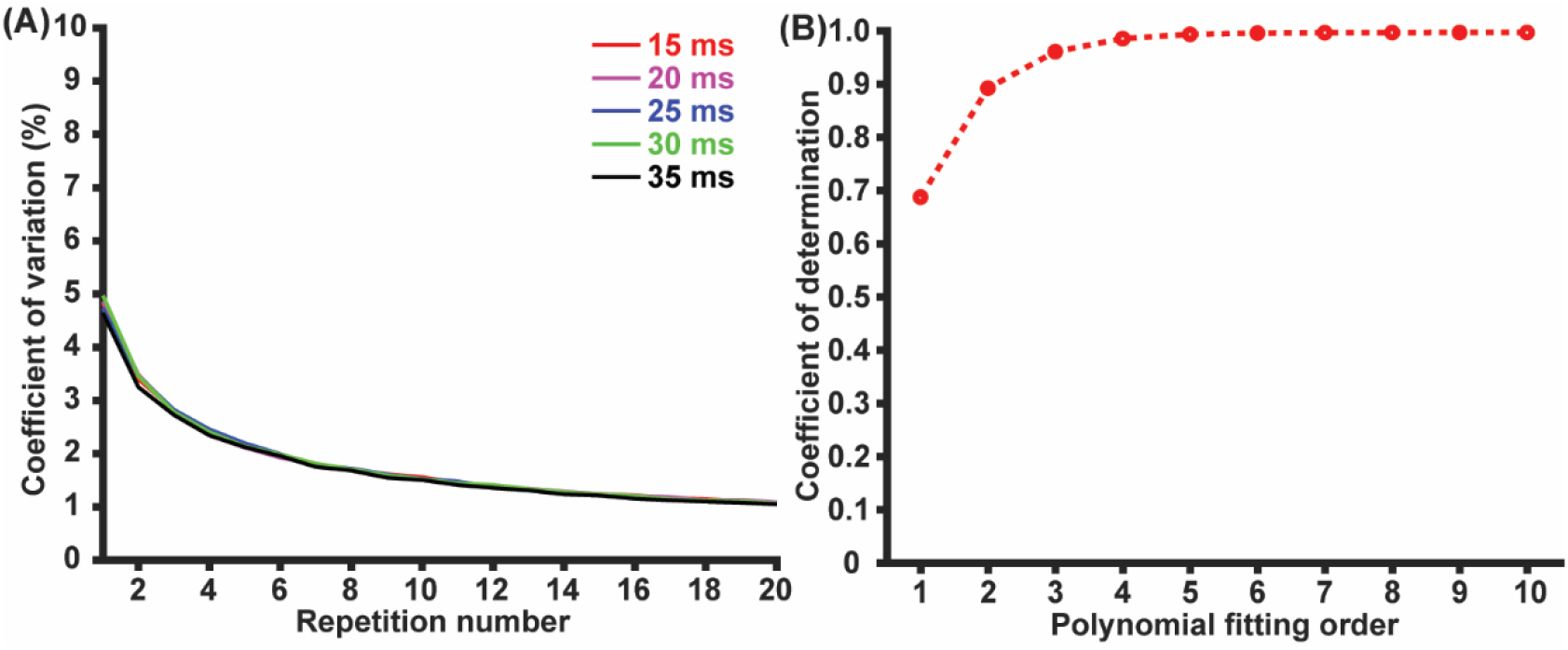
CoV as functions of repetition number (A) and coefficient of determination as a function of polynomial fitting order.

## Discussion

We have systematically investigated the influence of critical parameters in T_2_ mapping on the measurement reliability of transversal relaxation time. It was found that noise level is the most critical factor to affect the T_2_ measurement reliability, and TE range, TE number, and repetition number can be carefully determined to provide optimal performance at the same time cost.

Signal-to-noise ratio (SNR) at the normalized noise level of 2.2% is 33.2 dB. The SNR effect on measurement reliability will determine the highest spatial resolution of T_2_ mapping. If high spatial resolution is desired for comparing fine structures, more averages should be collected or advanced receiver coil (e.g., cryoprobe) can be employed. Monte Carlo simulations similar to those performed in the current study can be performed to determine the best spatial resolution at a given scan duration and hardware configuration.

The nonlinear dependence of measurement reliability on TE range suggests that proper coverage over the decaying signal is critical for T_2_ mapping. When TE range was narrow, the measured signals covered a small dynamic range close to the maximal signal (when TE = 0 ms). When TE range was too large, most of the measured signals were close to zero, leaving majority of the signal dynamic range underrepresented. According to our simulation, optimal TE range is 1.857 times the T_2_ to be measured. In practical applications, a prior knowledge on the T_2_ values is often unavailable and a same TE range will be used to measure different T_2_ values. If an additional 10% of the CoV at the TE range of 1.857T_2_ is allowed, 1.255T_2_ ≤ TE range ≤ 3.315T_2_ is acceptable. If the dynamic range of T_2_ to be measured is too broad, it is doable to utilize a large TE range beyond 3.315T_2_ together with increased TE numbers.

Both TE number and TE repetition can be increased to enhance measurement reliability. Note that the underlying principles are different. A larger number of TE indicates denser coverage over the dynamic range of signals. A larger number of TE repetition indicates more averages to improve the SNR. Both methods will lead to longer scan time.

Transversal relaxation time in MRI is sensitive to both structural and physiological changes (*7,15,28*). Physiological MRI is an active subfield with fruitful technological advancement and clinical translations (*29-34*). Measurement of brain metabolism has been a niche market of positron emission tomography (PET) (*35, 36*), which requires an on-site cyclotron and invasive procedures. Technical developments of MRI in the past decades have allowed the non-invasive evaluation on brain metabolism (*21,22,37-39*). Such MRI techniques were performed to study the metabolic dysfunctions in normal aging and different neurological diseases (*40-44*). Enhanced measurement reliability will promote the utilization of T_2_-fitting based physiological MRI techniques in future pathophysiological studies (*45, 46*).

Findings in the current study should be interpreted with limitations. Different experimental parameters were simulated separately with the assumption that there was no interplay. Note that the simulated data with simultaneous parametric changes will be hyperdimensional and difficult to interpret. Projection to a lower dimension to focus on a specific parameter will in principle erase interplays. Therefore, if the interplay between certain parameters are of interest, simulations can be designed accordingly and conducted.

## Conclusion

Good reliability with ≤5.0% variation can be achieved when the normalized noise level is below 2.2% (equivalent to the SNR of 33.2 dB). Optimal TE range for measuring T_2_ is related to the T_2_ under evaluation. TE number and repetition number can be increased to reduce measurement variations.

## Acknowledgements

The author would like to thank Dr. Zhiliang Wei for proofreading and editing.

